# Top-DTI: Integrating Topological Deep Learning and Large Language Models for Drug Target Interaction Prediction

**DOI:** 10.1101/2025.02.07.637146

**Authors:** Muhammed Talo, Serdar Bozdag

**Affiliations:** Department of Computer Science and Engineering, University of North Texas, Denton, TX 76207, USA; BioDiscovery Institute, University of North Texas, Denton, TX 76207, USA; Center for Computational Life Sciences, University of North Texas, Denton, TX 76207, USA; Department of Mathematics, University of North Texas, Denton, TX 76207, USA

**Keywords:** Topological Data Analysis, Large Language Models, Drug Target Interaction, Computational Drug Discovery

## Abstract

**Motivation:** The accurate prediction of drug–target interactions (DTI) is a crucial step in drug discovery, providing a foundation for identifying novel therapeutics. Traditional drug development is both costly and time-consuming, often spanning over a decade. Computational approaches help narrow the pool of compound candidates, offering significant starting points for experimental validation. In this study, we propose Top-DTI framework for predicting DTI by integrating topological data analysis (TDA) with large language models (LLMs). Top-DTI leverages persistent homology to extract topological features from protein contact maps and drug molecular images. Simultaneously, protein and drug LLMs generate semantically rich embeddings that capture sequential and contextual information from protein sequences and drug SMILES strings. By combining these complementary features, Top-DTI enhances predictive performance and robustness.

**Results:** Experimental results on the public BioSNAP and Human DTI benchmark datasets demonstrate that the proposed Top-DTI model outperforms state-of-the-art approaches across multiple evaluation metrics, including AUROC, AUPRC, sensitivity, and specificity. Furthermore, the Top-DTI model achieves superior performance in the challenging cold-split scenario, where the test and validation sets contain drugs or targets absent from the training set. This setting simulates real-world scenarios and highlights the robustness of the model. Notably, incorporating topological features alongside LLM embeddings significantly improves predictive performance, underscoring the value of integrating structural and sequence-based representations.

**Availability:** The data and source code of Top-DTI is available at https://github.com/bozdaglab/Top_DTI under Creative Commons Attribution Non Commercial 4.0 International Public License.

## Introduction

Drug development is a costly and long-term process because of the expensive and labor-intensive nature of experimental assays. The approval process for a new drug in clinical practice generally takes 10 to 15 years, with associated costs ranging between $161 million and $4.54 billion (1, 2). The drug development process encounters significant failure rates due to safety concerns, lack of efficacy, and the constraints of traditional methods such as high-throughput screening, which is challenged by the complex nature of evaluating extensive compound libraries (3). To overcome these problems, computational methods that improve and automate traditional methodologies have become an essential resource in drug discovery.

Drug target interaction (DTI) prediction is a crucial component of drug discovery research to identify target proteins of a drug molecule. Docking simulations, ligand-based methods, and machine learning approaches are the three fundamental types of computational techniques for DTI prediction. A docking-based approach uses 3D structures of target proteins and drug molecules with simulations to find binding sites. However, this method has limitations related to their laborious processes and difficulties in modeling complex proteins (4). Ligand-based models, which involve the comparison of novel ligands to known protein ligands, exhibit poor results when the number of known ligands is limited. Ligand-based and docking simulation methods yield promising predictive results; however, their reliance on high-quality data for drug molecules and proteins restricts their applicability in DTI prediction tasks (5, 6).

The methodologies used in machine learning for DTI prediction can be categorized into two groups: similarity- and feature vector-based approaches. Similarity-based methods are based on the assumption that similar drugs have a tendency to interact with similar targets. These methods employ a variety of similarity metrics based on, e.g., chemical, ligand, expression, and side-effect data to predict DTIs. Feature vector-based methodologies represent drugs and targets as feature vectors, which make it easier to capture complex interactions and connections among distant molecular components. Drugs are usually represented using multiple techniques, including the Simplified Molecular Input Line Entry System (SMILES), molecular fingerprints, two-dimensional structural representations, learned embeddings, and molecular graphs. Proteins are represented by sequence-, structural-, and network-based feature vectors. Network-based representations, like protein-protein interaction networks and knowledge graphs, combine various sources of data to enhance feature extraction. Machine learning models are trained using these representations to predict DTIs.

Deep learning techniques have significantly enhanced DTI prediction by automatically extracting high-dimensional features and modeling non-linear relationships between drugs and targets. These methods usually encode drug and protein structures separately, concatenate their learned representations, and then use them as input for a classifier. For instance, DeepDTA (7) uses convolutional neural networks (CNNs) to extract features from protein sequences and drug SMILES strings for prediction of binding affinity. Similarly, DeepConv-DTI (8) employes a CNN model to predict DTI using protein sequences and Morgan fingerprints of drugs. Hybrid deep learning approaches that combine LSTM and CNN architectures, as well as models that integrate diverse features, such as protein sequence, structure, and drug chemical properties, have been used to improve the accuracy of DTI prediction (9, 10).

Graph Neural Networks (GNN) are also capable of accurately predicting DTI by learning intricate relationships of drugs and targets within networks. For example, GraphDTA (11) and MGNDTI (12) model drugs as molecular graphs and employ GNN for predicting drug-target affinity and DTI. The GSL-DTI model incorporates heterogeneous networks using meta-path-based graph convolution to learn drug and protein representations for DTI prediction (13).

Attention-based models have received great interest in this domain. DrugBAN (14) integrates a bilinear attention network to comprehend local interactions between drugs and targets. MolTrans (15) utilizes a self-attention mechanism to change structural embeddings. HyperAttentionDTI (16) employs the attention mechanism on the feature matrices, assigning an attention vector to each amino acid. CoaDTI (17) models the interaction information from the drug and protein modalities using a co-attention mechanism.

Large language models (LLMs) have emerged as powerful tools in deep learning, utilizing vast amounts of unlabeled data through self-supervised learning. For downstream tasks, LLMs trained on protein sequences and drug SMILES string representations provide informative and contextually rich features. Protein sequence LLMs such as ProtT5 (18), ESM2 (19), and ANKH (20) models are trained using over millions of amino acid sequences. Similarly, drug-related LLMs such as MoLFormer (21), ChemBERTa (22), and ChemGPT (23) have shown significant effectiveness in capturing chemical characteristics. In recent years, pre-trained models have been employed for DTI prediction.The ConPLex model (24) utilizes a pre-trained protein language model to acquire protein representations using the contrastive learning method, aligning proteins and drugs in a common latent space. Kang et al. (25) employed the pre-trained ChemBERTa model for chemical compounds and ProBERT for target proteins to predict DTIs. The DrugLAMP model (26) combines molecular graph and protein sequence features derived from protein language models for DTI prediction.

Despite advances in DTI prediction through similarity-based, graph-based, and deep learning approaches, these methods mainly rely on features such as chemical descriptors, protein sequences, and graph-based embeddings. However, they often overlook topological components and structural data, which are also critical to capture interactions between drugs and targets.

Topological Deep Learning (TDL) is an emerging research area that combines the concepts of Topological Data Analysis (TDA) with current machine learning methodologies (27). TDA utilizes algebraic topology to uncover fundamental structure of high-dimensional datasets by examining their topological characteristics, such as connectivity, loops, and voids. Persistent homology, a fundamental technique in TDA, provides a robust and noise-resistant framework for capturing and analyzing multi-scale topological patterns in data. The collaboration between TDA and deep learning allows models to extract and leverage higher-order structural information that is frequently neglected by traditional methods (28, 29).

There has been a growing demand for utilizing TDA to tackle complex and high-dimensional problems across multiple domains. For example, in biomedical imaging, TDL has significantly improved histopathological cancer detection (30) and biological image segmentation (31).In genomics, TDL has facilitated identification of intricate patterns within genetic data, providing insights into genetic diseases (32). In drug discovery, ToDD (27) uses multiparameter persistence homology, enhancing virtual screening performance by incorporating domain-specific chemical features.

In this study, we propose a novel computational framework called Top-DTI for DTI prediction by integrating embeddings learned from TDA and LLMs. These embeddings are dynamically fused and further refined within a GNN, using the connectivity of the DTI graph. Our model leverages two key input features for both drugs and targets: embeddings from pre-trained drug and protein LLMs, and topological features derived through TDA on drug molecular images and protein contact maps.

To effectively combine these complementary features, we designed a feature fusion module that dynamically integrates TDA and LLM embeddings by assessing their relative importance during training. This fusion enhances the model’s ability to utilize both sequence-based and topological information. The integrated embeddings were subsequently processed through a heterogeneous GNN, which models the relationships between drugs and targets. This approach enables the identification of complex interaction patterns while efficiently leveraging both network topology and the fused feature representations.

The proposed Top-DTI model was evaluated on publicly available BioSNAP (33) and Human (34, 35) benchmark datasets. Experimental results demonstrate that Top-DTI consistently outperforms state-of-the-art DTI prediction models, highlighting its effectiveness and robustness. The main contributions of this work are as follows:

- We present a DTI prediction framework that utilizes TDA through the cubical persistence features derived from 2D drug molecule images and protein contact maps.
- The proposed framework uses MoLFormer and ProtT5 LLMs to extract 1D embeddings from drug SMILES strings and target protein sequences, respectively.
- We design a feature fusion module that dynamically fuses LLM and TDA embeddings for drugs and targets during training.
- We demonstrate the effectiveness of the Top-DTI model by conducting a comprehensive assessment on public benchmark datasets, obtaining superior results compared to the state-of-the-art methodologies.
- We examine the robustness of the proposed model in a challenging cold-split scenario, using unseen drugs and targets in the test set, imitating real-world situations.

Top-DTI provides a robust and efficient framework for DTI prediction by integrating complementary feature representations, thereby advancing computational drug discovery.

## Materials and methods

### Datasets

To evaluate Top-DTI, we used public BioSNAP and Human benchmark datasets, along with their variants, including BioSNAP unseen drugs, BioSNAP unseen targets, and Human cold datasets, which were obtained from (26) and (24). The BioSNAP dataset is derived from the DrugBank database and includes genes that are targeted by drugs available in the US market. The dataset is balanced, having verified 13,830 positive interactions and 13,634 negative interactions randomly sampled from non-interacting drug-target pairs. The Human dataset contains 2,633 positive interactions and 3,364 highly accurate negative interactions obtained by an *in silico* screening process. The datasets were divided into training, validation, and test sets and included protein sequences, drug SMILES strings, and interaction information for all pairs of drugs and targets. To ensure a fair comparison with other methods, we used the same training and test splits as used in the prior work. The statistical details of the benchmark datasets are given in Table 1.

**Table 1.** Statistics of benchmark datasets.

| Dataset | Drugs | Targets | Train | Validation | Test |
| --- | --- | --- | --- | --- | --- |
| BioSNAP Random | 4505 | 2181 | 9684 / 9540 | 1398 / 1349 | 2748 / 2745 |
| BioSNAP Unseen Drugs | 4505 | 2181 | 9531 / 9607 | 1383 / 1353 | 2916 / 2674 |
| BioSNAP Unseen Targets | 4505 | 2181 | 9872 / 9489 | 1382 / 1386 | 2576 / 2759 |
| Human Random | 2726 | 2001 | 1846 / 2351 | 251 / 349 | 536 / 664 |
| Human Cold | 1813 | 1503 | 1815 / 1638 | 59 / 96 | 121 / 190 |

We present the number of unique drugs and targets in each dataset. The number pairs in the train, validation, and test columns are reported as positive/negative, where positive indicates known interactions and negative indicates non-interacting drug-target pairs.

We have created two-dimensional representations of drug molecular structures and protein contact maps to capture the structural features of drugs and protein targets. The images of drug molecules were generated from their SMILES representations utilizing the RDKit library (36). To create protein contact maps, we used a transformer-based contact prediction model (37). The contact maps were created using the self-attention maps of the transformer model.

### The proposed method

The proposed framework called Top-DTI for DTI prediction integrates features derived from TDA and LLMs. Initially, TDA methods were utilized to derive topological features from molecular images of drugs and protein contact maps. Additionally, ProtT5 and MoL-Former LLMs were used to extract embeddings from protein sequences and drug SMILES strings, respectively. Then, the TDA and LLM embeddings were combined through a learnable fusion mechanism that dynamically balances the contributions of topological and sequence-based features. Afterwards, these integrated representations were fed to a heterogeneous GNN to learn relational information from the DTI network. Finally, embeddings learned from GNN were utilized to train a multilayer perceptron (MLP) classifier to predict DTIs. Figure 1 illustrates the Top-DTI framework.

**Fig. 1.**
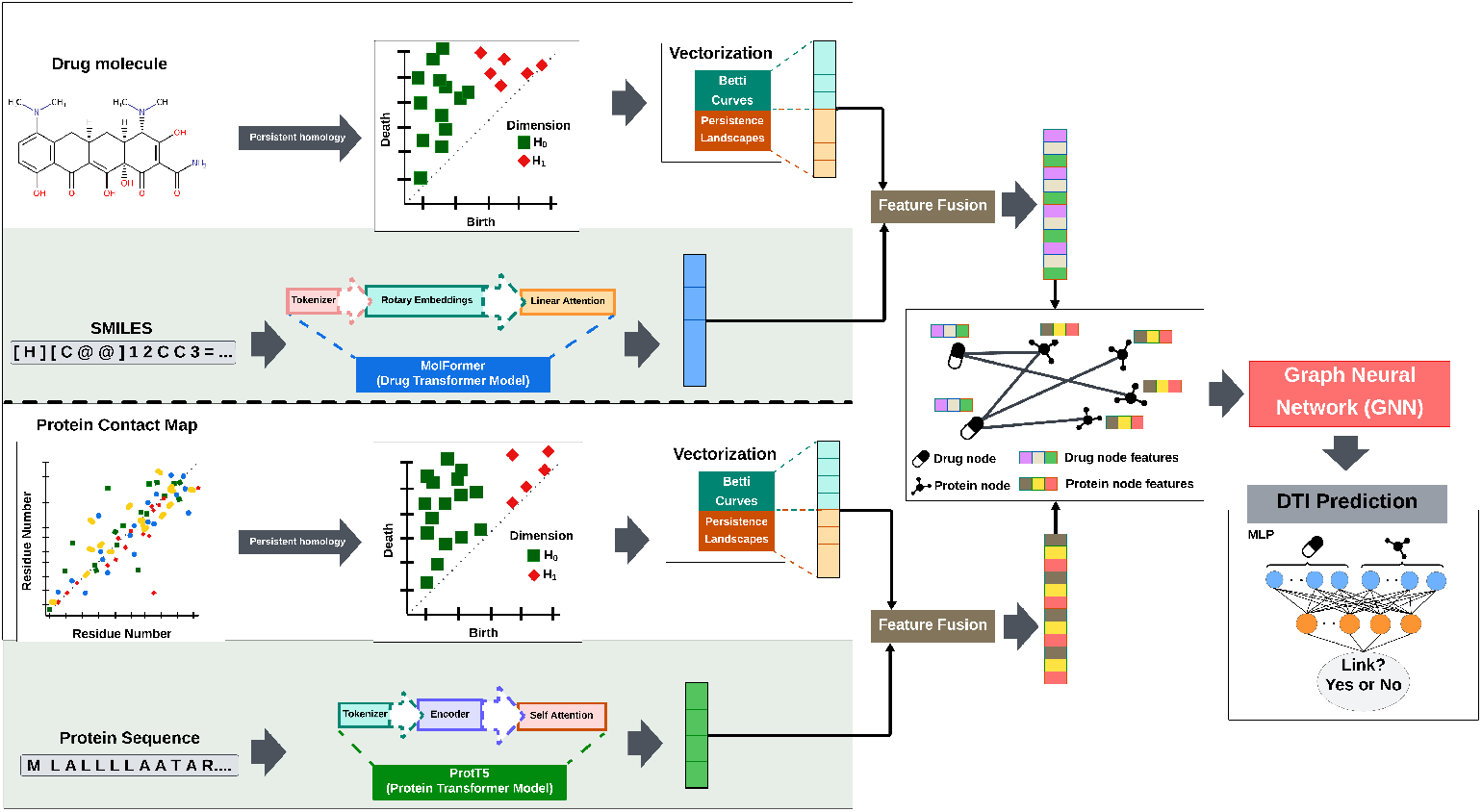
The Top-DTI model architecture includes vectorization of drug and target images using Betti curves and persistence landscapes. MoLFormer and ProtT5 transformer models are employed to extract sequential embeddings. Topological and sequential embeddings are combined using the *feature fusion* module, and the integrated representations are fed into a GNN. The embeddings obtained from GNN are concatenated and forwarded to a multilayer perceptron (MLP) to predict DTI.

### Topological feature extraction from drug molecular images and protein contact maps

In this study, we utilized persistent homology (PH) to extract topological features from two-dimensional molecular images of drugs and protein contact maps. PH is a method in TDA that analyzes structural information to capture topological features such as connected components and loops. These features are essential for capturing hidden shape patterns. The process of PH can be outlined in three steps. The first step, **filtration**, involves the construction of a nested sequence of cubical complexes from the images by monitoring the evolution of topological structures. The second step, **persistence diagrams**, records the *birth* (appearance) and *death* (disappearance) of topological features during the filtration process. The third step, **feature vectorization**, converts persistence diagrams into feature vectors using methodologies such as *Betti curves* and *persistence landscapes*.

Filtration is defined based on pixel values for images. For a given image *X* with size *r* × *s*, each pixel Δ_*ij*_ in the image has a value 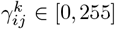, where *k* represents the selected color channel (grayscale, red, green, or blue). To construct the filtration, we choose a series of thresholds 0 = *t*_1_ *< t*_2_ *<* · · · *< t*_*N*_ = 255, where *N* is the number of thresholds. At each threshold *t*_*m*_, a binary image 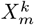 is generated as:

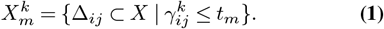

where, 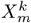 includes all pixel values less than or equal to *t*_*m*_ for the color channel *k*. At lower thresholds, only low-pixel-value regions are activated. As the threshold increases, more pixels are added, resulting in the formation of loops and larger connected structures. The result is a nested sequence of binary images (cubical complexes),

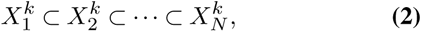

where additional pixels are “activated” as the threshold (pixel values) increases (28).

After the filtration, we compute persistence diagrams for dimensions *H*_0_ and *H*_1_. Here, *H*_0_ and *H*_1_ represents connected components and loops, respectively. Each persistence diagram consists of points (*b*_*i*_, *d*_*i*_), where *b*_*i*_ and *d*_*i*_ represents the birth and death times of a topological feature during the filtration, respectively. Features farther from the diagonal have longer lifespans and are considered significant, while features closer to the diagonal are typically noise.

Figure 2 illustrates this process with an example of a molecular image and its corresponding persistence diagram. In this figure, connected components (*H*_0_) are highlighted as green squares, and loops (*H*_1_) are represented by red diamonds. The connected components monitor distinct regions or components in the molecular structure, such as bonds between atoms. Loops (holes) correspond to cyclic substructures, such as rings in a molecular framework. A loop farther from the diagonal indicates persistent, significant cyclic structures. For example, four red diamonds positioned away from the diagonal represent the four rings in a given sample molecular image. In contrast, features near the diagonal have shorter lifespans and are more likely to represent noise or less significant variations.

**Fig. 2.**
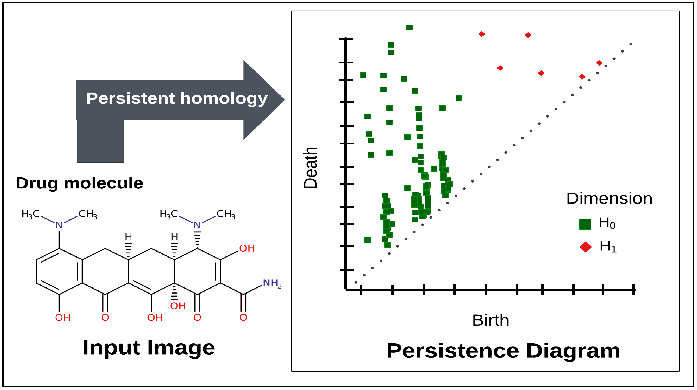
A sample of molecular image (left) and its corresponding persistence diagram (right). The green squares (*H*_0_) represent connected components, while the red diamonds (*H*_1_) represent loops.

After creating the persistence diagrams, we converted these diagrams into feature vectors by using vectorization methods, such as betti curves and persistence landscapes. Betti curves summarize the evolution of topological features across a filtration. For a given dimension *d*, the Betti number *β*_*d*_ represents the count of *d*-dimensional features in a topological space. In our case, *β*_0_ denotes the number of connected components, and *β*_1_ represents the number of loops in a binary image. Betti vectors are defined as:

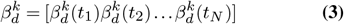

where *d* represents the dimension and *k* denotes the color channel. Using 50 thresholds (*t*_1_, *t*_2_, …, *t*_50_), and two dimensions (*d* = 0, 1), we computed Betti vectors for each dimension (*H*_0_ and *H*_1_). Therefore, we obtained a 100-dimensional vector for each color channel.

We utilized the persistence landscapes method alongside Betti curves to enhance the representation of structural patterns. Persistence landscapes directly use the lifespan of features. For each birth–death pair (*b, d*) in a persistence diagram, a piecewise-linear function *f*_(*b,d*)_ : ℝ → [0, ∞) is defined as:

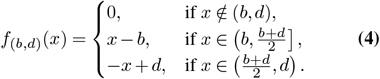

which creates a triangular shape over the interval (*b, d*), with a peak at 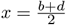. Persistence landscape (38) is defined as the sequence of functions *λ*_*l*_ : ℝ → [0, ∞), *l* = 1, 2, … where:

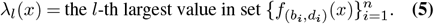

In Equation 5, *n* is the total number of birth-death pairs. We calculated persistence landscapes for the *H*_0_ and *H*_1_ dimensions using the first layer of persistent landscapes (the largest value, *λ*_1_). To achieve computational efficiency, we partitioned the filtration parameter range into 100 sampling points, referred to as bins. The feature vector for each color channel ends up being 200, as we performed an analysis on two homology dimensions.

As a result, Betti curves generated a 100-dimensional vector, and persistence landscapes contributed a 200-dimensional vector for each color channel, resulting in a total size of 300 dimensions. By concatenating the feature vectors across all four channels (grayscale, red, green, and blue), we obtained a 1200-dimensional feature vector. These representations were used as topological features in the TOP-DTI model.

### Sequence-based feature extraction using LLMs

We employed pre-trained LLMs to capture the sequence-based features of drugs and protein targets. We utilized the ProtT5 model for protein sequences and the MoLFormer architecture for drug representations. The ProtT5 model employs a transformer architecture featuring an encoder-decoder configuration and is trained on the UniRef50 database to predict masked amino acids with a masking ratio of 15%. We obtained 1024 dimensional protein target embeddings from the last hidden states of the model’s encoder. The resulting embeddings were averaged across sequence lengths to generate a fixed-size representation for each protein.

For drug molecules, we utilized MoLFormer, a transformer-based molecular LLM framework, to produce drug representations from chemical SMILES strings. MoLFormer was pre-trained on 1.1 billion unlabeled molecules sourced from the PubChem and ZINC databases (21). MoLFormer utilizes rotary position embeddings and an efficient linear attention mechanism to encode the spatial relationships of molecular structures. In the pre-training phase, 15% of the tokens in a SMILES string are masked for prediction. The pre-training task did not involve DTI prediction, thus there is no risk of data leakage. We have obtained 768 dimentional embeddings for drug molecules.

After the extraction of LLM embeddings and topological features of drugs and target proteins, two fully connected layers with ReLU activations were used to reduce their dimensionality and project them into a common 512-dimensional latent space to reduce computational complexity. These embeddings were used as high-dimensional sequence-based representations in our proposed Top-DTI framework.

### The integrated modeling of sequence and structural embeddings using dynamic feature fusion and GNN

In the Top-DTI framework, we implemented a *feature fusion* module for drugs and targets to efficiently fuse complementary information from sequence-based embeddings and topological features. During training, this module dynamically assigns weights to each feature type, facilitating the creation of enhanced embeddings. In particular, the sequence-based embeddings and topological features are concatenated, and the fully connected layer provides a weighting factor by sigmoid activation function (Equation 6).

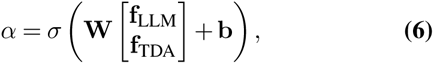

where *σ* is the sigmoid activation function, **f**_LLM_ ∈ ℝ^512^ denotes the sequence embedding vector of a drug or protein, **f**_TDA_ ∈ℝ^512^ denotes the topological feature vector of a drug or protein, **W** ∈ℝ^512*×*1024^ is a learnable weight matrix, and **b** ∈ℝ^512^ is a learnable bias vector. The sigmoid activation function ensures that the weight vector *α* ∈ℝ^512^ is in the range of [0, 1].

Using the weight vector *α*, the fused embedding **f** is computed as:

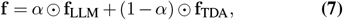

where ⊙ represents the element-wise multiplication. In Equation 7, the complement 1 − *α* is applied element-wise to adjust the contribution of topological features in the fusion process.

The fused features of drugs and targets were utilized as node features in a heterogeneous DTI graph. The nodes in this graph represent drugs and targets, while the edges indicate potential interactions. A two-layer GNN with SAGEConv architecture was applied to learn enriched embeddings for drugs and targets (39). Batch normalization and dropout layers were implemented at each layer to enhance generalization and mitigate overfitting.

After generating embeddings for drugs and targets using GNN, these embeddings were concatenated and used to train an MLP to predict DTIs.

### Model training and evaluation

We evaluated the performance of the Top-DTI model in two evaluation settings: *random-split* and *cold-split*. The random-split refers to the traditional method of dividing data into training, validation, and test sets, where drugs and targets can appear in both the training and test sets with different interactions. The random-split setting, while useful for preliminary evaluation, may not fully reflect the challenges of predicting interactions for unseen drugs or targets. In contrast, the cold-split setting ensures that any drug or target in the test set is not present in the training set. The cold-split scenario mimics real-world scenarios requiring the model to predict interactions for unseen drugs and targets. Therefore, the model must generalize the interaction beyond its training samples.

In the random-split setting, the interactions in the BioSNAP and Human datasets were split into 70% for training, 10% for validation, and 20% for testing by the dataset curators (Table 1). We used the same splits as the previous studies to have a fair comparison with state-of-the-art (SOTA) methods (33–35).

In the cold-split setting, Human dataset, prepared by Luo et al. (14), was designed to ensure that all drugs and proteins in the validation and test sets were excluded from the training set, thereby minimizing the risk of hidden data bias, as outlined in (35). **Unseen drugs** and **unseen targets** are variations of the BIOSNAP dataset in which drugs or targets in the test set are absent from any interactions in the training set. The unseen targets dataset was created by picking 20% of the targets from the entire dataset, including all interactions related to these proteins in the test set, guaranteeing that there is no target overlap between the training and test sets. The same methodology was applied to generate the unseen drugs dataset. The remaining dataset was further divided, allocating 7/8 of the interactions for training and 1/8 for validation (24).

The proposed Top-DTI model was trained using a binary cross-entropy loss function and Adam optimizer to update parameters. In order to mitigate overfitting, validation loss was monitored at each epoch, and an early stopping criterion was applied to terminate training if validation loss did not reduce for five consecutive epochs.

We used the area under the receiver operating characteristic curve (AUROC), the area under the precision-recall curve (AUPRC), sensitivity, and specificity as evaluation metrics. The results were reported as average performance over five independent runs, each initialized with a different random seed. All model outputs, including detailed performance metrics, were provided in Jupyter Notebooks to facilitate reproducibility. The source code and documentation are available at https://github.com/bozdaglab/Top_DTI.

## Results and discussion

To evaluate Top-DTI, we compared it against SOTA and baseline methods using the BioSNAP and Human benchmark datasets using both traditional random-split and more challening cold-split scenarios. We also performed an ablation study to examine the contribution of different components of Top-DTI. These analyses are summarized in the subsequent sections.

### Baseline and SOTA Methods

We evaluated the performance of Top-DTI model by comparing it with the following baseline and SOTA methods, including random forest (RF) and support vector machines (SVM), using the benchmark results reported in a most recent study by Luo et al. (26).

- DeepConv-DTI (8) employs a CNN to extract local residue patterns from raw protein sequences and combines these features with drug fingerprints.
- GraphDTA (11) model represents drugs as molecular graphs and utilizes a GNN to learn chemical features and interactions directly from the graph architecture.
- MolTrans (15) uses a frequent consecutive subsequence (FCS) mining algorithm to extract substructures from proteins and drugs and then encodes them using a transformer-based architecture to predict DTI.
- DrugBAN (14) integrates a GCN to encode local structures from drug molecular graphs with a CNN to encode protein sequences. Then, a bilinear attention network processes these encoded features for DTI prediction.
- DLM-DTI (40) adopts the ChemBERTa model for drug feature extraction and the ProtBERT transformer for protein representations. These features are concatenated and used to train an MLP architecture to predict DTI.
- Kang et al. (25) employ ChemBERTa and ProtBERT transformer-based models to encode drug molecules and protein sequences and implement prediction using a classifier.
- DrugLAMP (26) uses a multimodal framework that combines molecular graph and protein sequence features extracted from the ESM-2 and ChemBERTa-2 LLMs. Pocket-guided co-attention and paired multi-modal attention are applied to encode these features to predict DTI.

### Comparison of Top-DTI with baseline and SOTA methods

We evaluated the performance of the Top-DTI model by comparing it with the baseline and SOTA methods reported in a recent study by Luo et al. (26). In a standard random-split setting, the performance comparison of Top-DTI with SOTA and baseline methods on the BioSNAP dataset is presented in Table 2.

**Table 2.** Performance comparison of methods on the **BioSNAP** dataset. Sens: sensitivity, Spec: specificity. Best and second best result for each evaluation metric are shown in bold and underlined, respectively.

| Method | AUROC | AUPRC | Sens. | Spec. |
| --- | --- | --- | --- | --- |
| SVM | 0.862 | 0.864 | 0.711 | 0.841 |
| RF | 0.860 | 0.886 | 0.823 | 0.786 |
| DeepConv-DTI (8) | 0.886 | 0.890 | 0.760 | 0.851 |
| GraphDTA (11) | 0.887 | 0.890 | 0.745 | 0.854 |
| MolTrans (15) | 0.895 | 0.897 | 0.818 | 0.831 |
| Kang et al. (25) | 0.910 | 0.900 | <u>0.862</u> | 0.847 |
| DrugBAN (14) | 0.903 | 0.902 | 0.820 | 0.847 |
| DLM-DTI (40) | 0.914 | 0.914 | 0.848 | 0.844 |
| DrugLAMP (26) | <u>0.917</u> | <u>0.922</u> | 0.844 | <u>0.855</u> |
| **Top-DTI** | **0.939** | **0.941** | **0.866** | **0.857** |

Top-DTI demonstrated superior performance compared to existing SOTA methods, including Kang et al. (25), DLM-DTI, and DrugLAMP, which mainly depend on LLM-based drug and target embeddings. Specifically, Top-DTI demonstrated an improvement above 2% in both AUROC and AUPRC compared to DrugLAMP, the previous SOTA method. The sensitivity and specificity scores are balanced, which shows that the TOP-DTI model precisely detects both interacting and non-interacting drug-target pairs.

Table 3 presents the results of the performance comparison of Top-DTI with SOTA and baseline methods on the Human dataset in a random-split setting. Top-DTI shows superior performance compared to all approaches, obtaining the highest AUROC and AUPRC values.

**Table 3.** Performance comparison of methods on the **Human** dataset. Best and second best result for each evaluation metric are shown in bold and underlined, respectively.

| Method | AUROC | AUPRC |
| --- | --- | --- |
| SVM | 0.940 | 0.920 |
| RF | 0.952 | 0.953 |
| DeepConv-DTI | 0.980 | 0.981 |
| GraphDTA | 0.981 | 0.982 |
| MolTrans | 0.980 | 0.978 |
| DrugBAN | 0.982 | 0.980 |
| DrugLAMP | <u>0.985</u> | <u>0.983</u> |
| **Top-DTI** | **0.993** | **0.992** |

Notably, all methods demonstrate high scores for this dataset, with AUROC and AUPRC values greater than 92%. In (35), Chen et al. indicated that there could be biases in the Human dataset due to single-class ligands and algorithmically generated negative samples. The superior predictive performance of the models may be linked to this biased information. The cold-split setting resolves this issue by guaranteeing that test drugs and targets are completely removed from the training set. This design strategy requires all models to generate predictions without prior knowledge of specific drugs or targets (14). Table 4 presents the results for the Human Cold dataset in the cold-split setting.

**Table 4.**
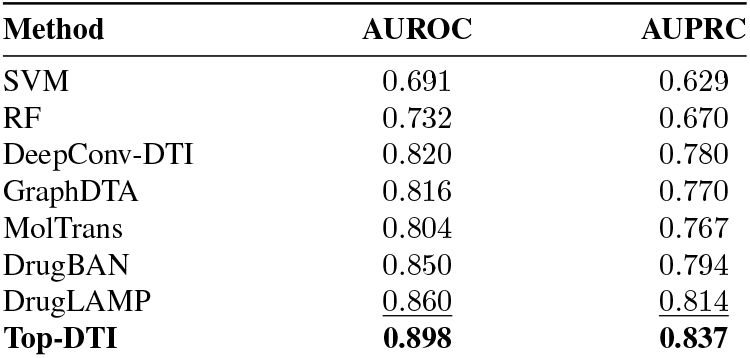
Performance comparison of methods on the **Human Cold** dataset. Best and second best result for each evaluation metric was shown in bold and underlined, respectively.

| Method | AUROC | AUPRC |
| --- | --- | --- |
| SVM | 0.691 | 0.629 |
| RF | 0.732 | 0.670 |
| DeepConv-DTI | 0.820 | 0.780 |
| GraphDTA | 0.816 | 0.770 |
| MolTrans | 0.804 | 0.767 |
| DrugBAN | 0.850 | 0.794 |
| DrugLAMP | <u>0.860</u> | <u>0.814</u> |
| **Top-DTI** | **0.898** | **0.837** |

As expected, all model performances decreased; however, Top-DTI still outperformed all the methods. In particular, compared to the previous best method, Top-DTI showed an improvement over 4.4% in AUROC and 2.8% in AUPRC.

### Ablation study

To evaluate the contributions of different components of Top-DTI, we performed ablation studies using the unseen drugs and unseen targets datasets. We compared the performance of Top-DTI with seven ablation models, each leveraging distinct combinations of topological and LLM features as follows:

- **Top_D + Top_T**: Integrates only topological features for both drugs and targets.
- **Top_D + LLM_T**: Integrates topological features of drugs with LLM embeddings of targets.
- **LLM_D + Top_T**: Integrates LLM embeddings of drugs with topological features of targets.
- **LLM_D + LLM_T**: Integrates only LLM embeddings for both drugs and targets.
- **Static Fusion (***α* = 0.5): Integrates LLM embeddings with topological features by assigning equal weight to both modalities with a fixed *α* value of 0.5.
- **Betti Fusion** : Integrates LLM embeddings with only Betti topological features using dynamic *α* values.
- **PL Fusion** : Integrates LLM embeddings with only Persistence Landscape topological features using dynamic *α* values.
- **Dynamic Fusion (proposed method)**: Integrates both LLM embeddings and topological features using both Betti and persistence landscape features along with dynamic *α* values.

The performance comparison of the Top-DTI model with different feature types for unseen drugs and unseen targets in BioSNAP dataset is given in Table 5. The results demonstrate that the model that included only LLM or topological features demonstrated the worst performance, underscoring the limitations of single feature types. Additionally, the integration of drug topological features with LLM embeddings in the unseen drug dataset improves model performance. Furthermore, the Static Fusion model, with an equal weight of 0.5 assigned to each data modality, outperforms single feature type methods, highlighting the value of integrated feature approaches. Both Betti Fusion and PL Fusion also surpass single feature type models and demonstrate competitive performance with the Static Fusion model. Finally, the Top-DTI model, which dynamically weights LLM and topological features while utilizing both Betti and persistence landscape embeddings, achieves the highest AUROC and AUPRC for both unseen drugs and targets. These results suggest that the adaptive integration of topological features with LLM embeddings further improves performance due to the complementary benefit of both representations.

**Table 5.** Performance of the Top-DTI model with **Feature Combinations** on Unseen Drug and Unseen Target Datasets.

| Features | Unseen Drug |  | Unseen Target |  |
| --- | --- | --- | --- | --- |
|  | AUROC | AUPRC | AUROC | AUPRC |
| Top_D + Top_T | 0.887 ± 0.003 | 0.907 ± 0.002 | 0.855 ± 0.006 | 0.866 ± 0.005 |
| Top_D + LLM_T | 0.899 ± 0.003 | 0.917 ± 0.002 | 0.898 ± 0.003 | 0.893 ± 0.005 |
| LLM_D + Top_T | 0.905 ± 0.003 | 0.919 ± 0.003 | 0.869 ± 0.007 | 0.870 ± 0.006 |
| LLM_D + LLM_T | 0.889 ± 0.004 | 0.905 ± 0.003 | 0.894 ± 0.003 | 0.889 ± 0.003 |
| Static Fusion ($\alpha = 0.5$) | 0.907 ± 0.005 | 0.921 ± 0.004 | <u>0.906 ± 0.004</u> | <u>0.904 ± 0.006</u> |
| Betti Fusion | 0.907 ± 0.002 | <u>0.922 ± 0.002</u> | 0.902 ± 0.002 | 0.898 ± 0.002 |
| PL Fusion | <u>0.908 ± 0.004</u> | 0.921 ± 0.004 | 0.903 ± 0.004 | 0.901 ± 0.003 |
| **Dynamic Fusion (Proposed)** | **0.911 ± 0.003** | **0.924 ± 0.002** | **0.907 ± 0.003** | **0.904 ± 0.003** |

In the final step of our analysis, we examined the dynamic weighting of the feature fusion module to understand how the Top-DTI model aligns the topological and LLM-based embeddings in the unseen target dataset. For this, we calculated the mean of the dynamically assigned weights (*α*) for both drugs and targets at each epoch during training. The weight vector *α* reflects the relative significance of drug and target feature types throughout the training process, as shown in Figure 3.

**Fig. 3.**
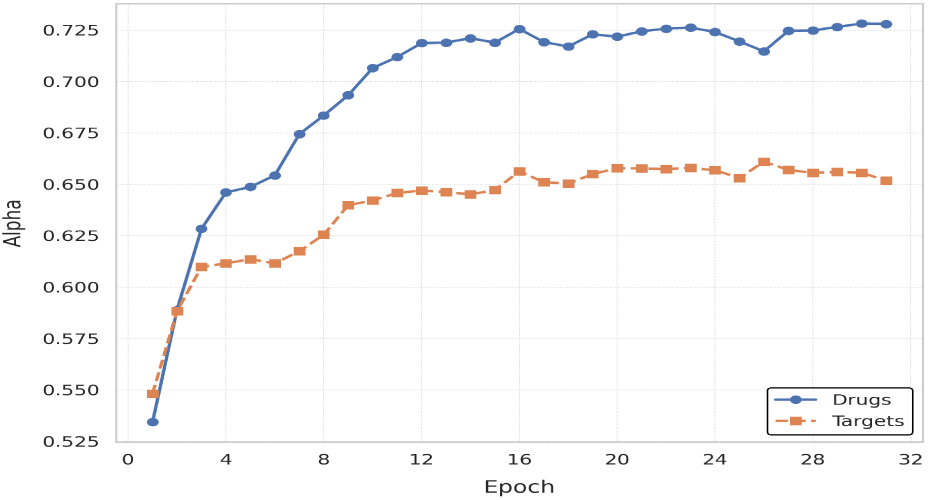
Mean alpha values for drugs and targets during training.

The TOP-DTI model assigns greater weights to LLM-based embeddings, 72% for drugs, and 64% for targets. The ablation study also showed that using only LLM-based embeddings performed better than using only topological features (Table 5). These findings demonstrate how each feature type contributes to the TOP-DTI model with the semantic richness of sequential representations and the complementary role of topological features.

## Conclusion

Drug-target interaction (DTI) prediction plays a critical role in the drug development process, as it can improve efficiency while reducing time and costs. In this study, we investigate the integration of topological data analysis (TDA) with large language models (LLMs) for the prediction of DTI. We used TDA to extract topological features from two-dimensional representations of drugs and protein targets and utilized LLMs to encode sequence-level information of both drugs and proteins. The proposed model, Top-DTI, significantly enhanced DTI prediction by integrating topological features with sequential embeddings using a dynamic feature fusion module. Future research could further improve DTI prediction by integrating multi-omic data, such as gene expression and proteomics for targets and molecular features for drugs.

## ACKNOWLEDGEMENTS

This work was supported by the National Institute of General Medical Sciences of the National Institutes of Health under Award Number R35GM133657 and the startup funds from the University of North Texas.

